# Stress anisotropy in 3D active curved structures

**DOI:** 10.1101/2025.04.13.648577

**Authors:** Yuting Lou, Sophie Theis, Jean-Francois Rupprecht, Timothy E Saunders, Tetsuya Hiraiwa

## Abstract

Layers composed of lateral connections prevail in biological systems from subcellular membranes to epithelial sheets. This work presents a continuum framework to describe the effects of mechanical forces within a 3D curved layer with supporting lateral mesh. We provide detailed discussion on the emergence of stress anisotropy as a function of depth in different curvature settings, building on Lou et al. Phys. Rev. Lett. **130**, 108401 (2023). We principally consider an epithelial monolayer to explain how the interplay between layer curvature and cell mechanics determines the stress anisotropy. We show that this can lead to irregular cellular shapes in 3D, including scutoid-like geometries. Our framework is general, and can be extended to a diverse set of biologically relevant systems.

## I. INTRODUCTION

How do complex forms emerge within active systems, such as living multicellular organisms [1]? There has been substantial work on understanding how flat tissues develop shape [2], as exemplified by the *Drosophila* wing disc [3–5]. Yet, most tissues and organs are complex, 3D structures [6–8]. Recent work has revealed that active systems in 3D can take on a diverse array of cell shapes [9–11]. Understanding when, where and how particular cell geometries emerge within active systems remains a challenge.

The surface of biological tissues is typically formed by a layer of confluent and tightly bound cells, known as an epithelium [12]. Epithelia provide an important barrier function, but also play roles in signaling and other cell processes [13]. Imaging of epithelial cells revealed that they generally have a hexagonal structure, consistent with predictions based on topological constraints [14]. Nevertheless, biological tissues undergo active events such as cell division and death (apoptosis) that alter cell packing within tissues [14, 15]. One particularly well characterized topological change within epithelia are T1-transitions at the apical surface of the tissue, whereby a group of four cells rearrange to alter the cell neighbors [16]. If these rearrangements are directed (*e*.*g*., by there being orientated actomyosin contractility), then such T1-transitions can lead to large-scale tissue deformations, such as in *Drosophila* germband elongation [16, 17], and frequent T1-transitions can effectively fluidize biological tissues [3, 18].

Recently, partly due to imaging advances, the full 3D shape of epithelia has become accessible. With such data, it has become apparent that the apical and basal surfaces do not necessarily behave concomitantly [19–22]. Indeed, the active T1-transitions observed during *Drosophila* germband elongation differ in their dynamics and mechanism between the apical and basal surfaces [19]. Even in tissues that were close to equilibrium, cell neighbors exchange between apical and basal surfaces; there were static apical-to-basal T1-transitions between the two surfaces [23–25]. We henceforth referred to these as AB-T1 transitions, to distinguish them from the dynamic T1-transitions discussed in the previous paragraph. Such cell rearrangements have been observed in highly curved tissues [24–26], tissues with high curvature anisotropy [23], tissues with significant rate of division, apoptosis or extrusion [27–29] and in foam models [30, 31].

Models have been proposed for the formation of AB-T1 transitions driven by topological constraints [23, 24, 32, 33]. However, these models consider only specific scenarios. Here, buiding on [33], we present detailed derivations of equations and discussions of a two-elastic-shell model for curved epithelia and the formalism is generalizable to multilayer systems with lateral compartments. We first outline a measure for the AB-T1 transition tendency and verify this through vertex simulations. We demonstrate that this measure is sensitive to cell plasticity. We demonstrate how such behavior impacts the cell shape in 3D. We then consider different biophysically relevant conditions. For simple analytics, we consider axisymmetric geometry, though the methods are applicable to arbitrary geometry. In the case where hydrostatic load dominates, we show that the rate of AB-T1 transitions increases with curvature anisotropy, which is maximal in a region such as the trunk of a prolate ellipsoid. We next apply our formalism to a scenario when tilt of cell lateral surfaces is achieved and show that AB-T1 transitions can also occur where high curvature anisotropy is induced by the onset of cell tilt. Overall, this work, in partnership with Ref. [33], builds a detailed theoretical understanding for how cell shape within confluent tissues is determined by geometric constraints in 3D.

## II. DEVIATORIC STRESS PREDICTS ONSET OF AB-T1S IN A VERTEX MODEL

In Ref. [33], we introduced a dimensionless measure for the tendency for AB-T1 transitions, *γ*, and demonstrated that

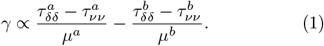

where *τ* with sub- and superscripts is a component of a tensor describing the tensions in-plane at either the apical (*a* superscript) or basal (*b* superscript) surfaces, with *δ* and *ν* denoting orthogonal in-plane coordinates. Positive *τ* indicates tension and negative *τ* compression. The denominator *µ* denotes the stiffness on each surface. The physical meaning of *γ* is the difference in the strain caused by anisotropic tensions between two layers.

To show how *γ* defined in Eq. 1 indicates the incidence of irregular cellular rearrangement, we performed vertex model simulations for an epithelial monolayer, which is a typical bilayer system with lateral compartments. We have outlined the results of such simulations in [33]. Here, we expand on these results to more thoroughly explore how the cell and tissue properties impact the relationship between *γ* and the tendency for AB-T1s to occur.

## A. Method

The simulations were performed on the Tyssue software [34]. We simulated apical and basal layers, connected by lateral bonds (Fig. 1a, left) with an energy function :

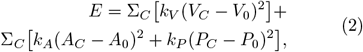

where *V*_0_, *A*_0_ and *P*_0_ are the preferred volume of the cell, preferred area and preferred perimeter of the apical/basal face respectively; *k*_*V*_, *k*_*A*_ and *k*_*P*_ are the elasticity strength parameters, respectively.

**FIG. 1.**
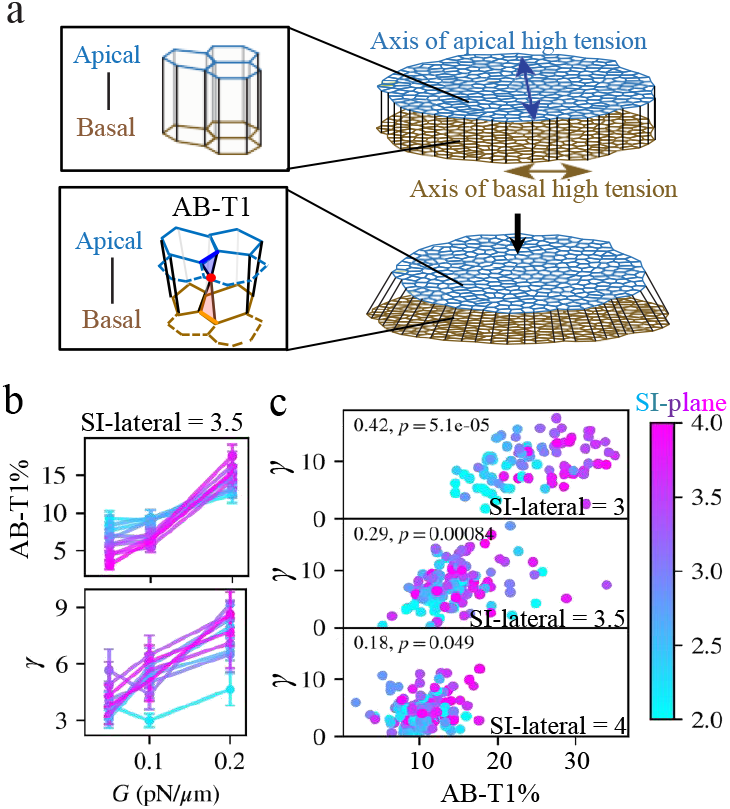
Vertex model simulations in 3D. (a) 3D simulation setting with two external stresses along two orthogonal axes. (b) The percentage of oriented AB-T1 transitions, AB-T1% (defined as the count of AB-T1 events normalized by the total cell number with the stochastic residue removed) and a quantity describing cell deviatoric deformation, *γ*, grow with anisotropic stress magnitude *G* for lateral shape index (SI-lateral) 3.5, under varying in-plane shape index (SI-plane). Color scheme is the same as in (c). Error bars represents standard error, from 10-12 simulation repetitions. (c) Comparing the AB-T1 transition percentage with *γ* under *G* = 0.2pN/*µ*m. Pearson R-correlation is shown in the left-top corner of each panel. Panels from top to bottom correspond to different values of lateral shape index (SI-lateral) and the color of dots correspond to the shape index of the cells at the apical/basal plane (SI-plane).

In order to create an anisotropy of tension in the tissue, we add a term of line tension in Eq. 2.

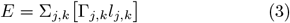

*l*_*j,k*_ is the length of the edge between vertices *j* and *k*, and Γ_*j,k*_ is a line tension specified as :

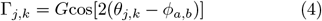

where *G* is the the amplitude, *θ*_*j,k*_ is the edge angle and *ϕ*_*a,b*_ is the angle of anisotropy for apical or basal surface.

Line tension will be maximum for edges parallel to the angle *ϕ* and minimum for edges perpendicular to *ϕ*. When edge length reaches a minimum threshold *l*_*j,k*_ *< l*_*t*_, a T1 transition is performed, leading to different cell neighbors between apical and basal sides of the cell, the left bottom panel in Fig. 1a, (*i*.*e*. the AB-T1 transition), following the implementation in [35].

By setting *ϕ* _*a*_ = *π/*2 and *ϕ* _*b*_ = 0, we can generate a deviatoric stress across the tissue depth (Fig. 1a, right) [33]. To quantify the adaptation of tissue to this stress, we characterized the number of cells that have differences in neighbors between apical and basal surfaces at the end of the simulation. This measure gives us a readout of emergent AB-T1 transitions as a result of the applied stress, since the simulation is initiated with identical neighboring configuration between apical and basal sides.

We previously showed in [33] that the number of AB-T1 transitions has a non-zero residue if no external stress is applied, and these transition present stochastic orientations. This is because of the system being initiated in a off-eqauilibrium state. Yet, with increase in the external stresses, AB-T1 transitions along the stress orientation occurs. We have introduced the quantity *γ* to describe this oriented deformation along two external stresses. *γ* can be represented by the normalized difference between the two eigenvalues of a statistical tensor that describes the mean orientation of AB-T1. Here, we briefly recall the definition of this statistical tensor. An AB-T1 implies four cells with two cells (here labeled 1 and 2) sharing a common junction on the basal side but not on the apical side and two other cells (here labeled 3 and 4) sharing a common junction on the apical side but not on the basal side. For each AB-T1, we define the matrix *m*_*a*_ (with *a* for apical) based on the pairwise distance between the cell centers 1 and 2 on the apical side (resp. 3 and 4 on the basal side), according to

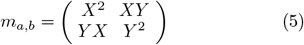

where 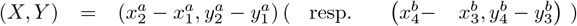, with 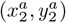 being the center of the face of the cell index 2 (resp. 4) within the AB-T1 quadruplet on the apical (resp. basal) side. For each simulation, we estimate the tensor

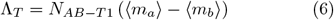

where *N*_*AB*−*T* 1_ is the total number of AB-T1s in this given simulation and the average is performed over all AB-T1 matrices defined in Eq. (5).

The two eigenvalues of ⟨Λ_*T*_⟩, denoted Λ_*x*_ and Λ_*y*_, describe how many edges appear, on average, on the apical side (with respect to the basal side) along the two orthogonal directions in a 2D plane. The tendency per cell for AB-T1 to occur along the external tensions (apical stretched along the *y* and basal stretched along the *x* axis) can then be calculated as:

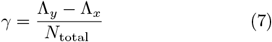

where *N*_total_ is the total number of cells then *γ* has dimension of area in the unit of *µ*m^2^; it is a static analogy to the tensor defined for the dynamic T1-transitions discussed in [36], where here the time argument is replaced by layer depth.

In our simulations, we have used the following parameter values: *l*_*t*_ = 0.1 *µ*m, *k*_*V*_ = 0.5u/*µ*m^6^, *k*_*A*_ = 1u/*µ*m^4^, *k*_*P*_ = 0.5u/*µ*m^2^, where *u* is the unit energy. The preferred area *A*_0_ = 1*µ*m^2^. The source code for the model is released under the GNU General Public Licence and is available on github at https://github.com/TimSaundersLab/CellPacking

## B. Results

Since *γ* measures the averaged appearance/disappearance of bonds from the basal to apical side along the direction of anisotropic stress, it has a physical meaning of deviatoric deformation across the layer. In our previous paper, we found that *γ* grows quasi-linearly with the external anisotropic stress magnitude *G* (Fig. S1B, [33]) and with the counted AB-T1 events as well (Fig. S1C, [33]). Here, we delve more deeply into the positive correlation between *γ, G* and the observed percentage of AB-T1 transitions, particularly under different cell shape pro<u>pe</u>rties.

The shape index (SI), defined by 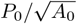, is a control parameter for the transition between fluid-to solid-like behaviors [18] and is straightforward to access experimentally. It determines the plastic deformability of cell boundaries, and larger shape index leads to higher energy dissipation through cell shape rearrangement, subsequently to more frequent intercellular shearing and to softer (more fluid-like) tissue [18]. In Fig.1b, we see that there is a relationship between increasing *G* and the increased oriented AB-T1 percentage and *γ*. This trend is weakened at very low shape factor. This corresponds to rigid “solid-like” cells, which are less able to reconfigure due to applied anisotropic stress.

As shown in Fig. 1c, there is a robust positive correlation between *γ* and the percentage of AB-T1 events along the anisotropic stresses, and the correlation is more significant for a lower shape index value on the lateral side (PearsonR correlation shown in the top left corner). Taking these results together, we see that *γ* serves as an effective proxy for the occurrence of AB-T1 events in a range of different physical conditions.

To conclude this section, we have established, on an athermal 3D cell-based model, that the average direction of AB-T1s follows the averaged deviatoric stress difference between the apical and basal sides and the positive relationship is robust against different plasticity in cellular arrangement (controlled by shape index of cell). In the next section, we turn to a continuum thin-shell model to detail how the deviatoric stress is modulated in the presence of an external curvature.

## III. CONTINUUM THIN-SHELL MODEL

Given the knowledge that the difference in anisotropic stresses between apical and basal surfaces leads to AB-T1s and irregular 3D shapes, we consider a minimal continuum model for a 3D epithelial layer and study the stress distributions therein. This model can also be extended to a more generalized layer system with lateral compartments. The modeled epithelium is composed of two elastic shell membranes connected by lateral interfaces Fig 2a, with the shell thickness *δ* much smaller than the depth of the layer *e*. Therefore, force balance is considered with merely the in-plane stresses within each shell, with the lateral forces as external forces exerted to the planes. Using the elastic shell model facilitates the analytical expressions of stress profiles in relation to curvature profiles under various assumptions that correspond to diverse biological conditions.

**FIG. 2.**
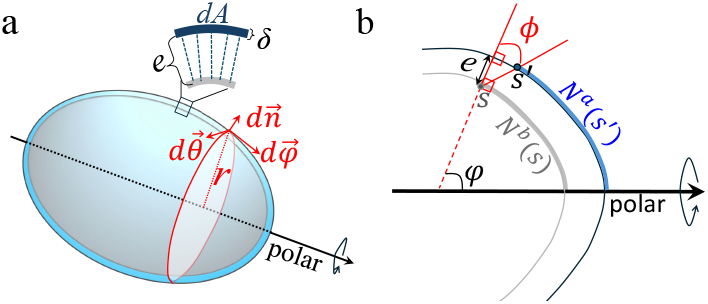
(a) A schematic two-layered elastic shell model with lateral support in 3D. (b) A meridional cross-section of an arbitrary axisymmetric shell with apical (navy) and basal (grey) layers. The apical layer is an outward projection of the basal layer along the normal direction at each local surface element with a distance *e*. The position *s*′ on the apical side corresponds to position *s* on the basal side in such a way that the cell number accumulated on the apical surface from the head to *s*′ equals the basal one accumulated from head to *s*; therefore, the angle between the vector from *s* to *s*′ (black bold line) and the surface normal direction (red dashed arrow) is the cell tilt angle *ϕ* describing the degree of cell tilt at the local surface.

### A. Force balance in axisymmetric shells

For a complete description of the model, we first include the force balance calculation here. This is also available in the Supplementary Material of Ref. [33].

We consider an object covered by an enclosed 3D surface with arbitrary curvature. Each point on the 3D surface is identified by the normal direction 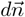, the meridional tangential direction 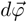 and the circumferential tan-gential direction 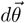 (Fig. 2a). In the following force-balance arguments, we limit our system to merely convex surfaces.

Assuming the system is near mechanical equilibrium and the surface is a purely elastic material, the stress tensor 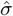 is balanced in three directions as:

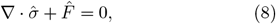

where 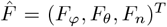 is the external body force imposed on an infinitesimal volume element. Hence, the full force balance equations in three dimensions read

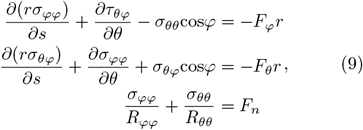

In a system with rotational symmetry around a polar axis, an infinitesimal volume element can be expressed as *dA·δ* = *rdθds*, with distance *r* to the polar axis, where *A* is the surface area, *δ* is the surface thickness, *s* is the arc length along the meridional direction and *rdθ* is the infinitesimal length along the circumferential direction. Let *R*_*φφ*_ and *R*_*θθ*_ be the radius of curvature along *ds* and *rdθ* respectively, and hence *R*_*φφ*_ = *ds/dφ* and *R*_*θθ*_ = *r/*sin*φ*. For the axisymmetric stresses, we ignore the derivatives of *∂θ* and set *F*_*θ*_ as zero. Then, from the second equation in Eq. 9, 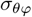 has to be zero, indicating that the torsion around the polar axis vanishes.

If the surface is sufficiently thin, *i*.*e. δ R*_*θθ*_, ≪ *R*_*φφ*_, across the layer thickness direction, all quantities can be taken as uniform so the transverse shear is neglected. Furthermore, bending stresses are also neglected at the discontinuity of displacement (usually at the apex of the object) due to the in-plane stresses [37–39]. With the assumptions of thin layers, we can first integrate Eq. 8 along the normal direction (over thickness) and arrive at a force balance only involving the in-plane tensions with unit N/m, which could be denoted as a tensor 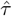, and neglecting derivatives along the normal directions. We can subsequently obtain the force balance equations in terms of the components of 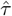:

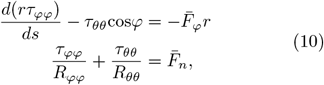

where 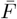 is the body force integrated over thickness, with units N/m^2^.

From Eqs. 10, we obtain a differential equation for *τ*_*ϕϕ*_

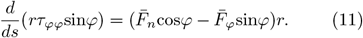

Integrating Eq. 11 over *ds* and multiplying a numerical factor 2*π* to the both sides, we arrive at:

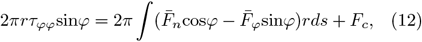

where the left-hand side is the total amount of force parallel to the polar axis at a latitudinal cross-section of the shell positioned with arc length *s* and the meridional angle *ϕ* (see Fig. 2b). This force is balanced by the summation of distributed load across the surface and a concentrated force *F*_*c*_ at the apex *s* = 0. Alternatively, this integral could be expressed by a definite integral from *s* = 0 to *s* = *s*(*ϕ*).

Since there is no concentrated force at the apex, *F*_*c*_ = 0. As the unit of 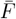 is N/m^2^, we re-denote the two external forces as the loading per unit area, the tangential one *σ*_*T*_ and the normal one *σ*_*N*_, such that 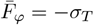 (with a minus sign so that *σ*_*T*_ *>* 0 points towards the apex) and 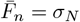 (positive pointing to the outward), respectively.

Therefore, the two force balance equations in relation to the curvature becomes

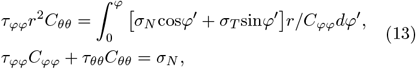

where *C*_*φφ*_ = 1*/R*_*φφ*_ and *C*_*θθ*_ = 1*/R*_*θθ*_ are the principal curvatures along meridional and circumferential directions.

### B. In-depth deviatoric strain

We next elaborate the rationale behind the derivation of the measure for AB-T1 transition tendency *γ* (Eq. 1). As shown in Sec. II, the number of AB-T1 events in our vertex simulations (Eq. 1) increased with the difference in the deviatoric stresses between the tissue layers (Fig. 1b-c, also see Fig.1 in [33]). In addition, the emergence of such AB-T1 transitions could be influenced by preferred cell shapes in the apical/basal and lateral planes. Motivated by these observations, we propose that the dimensionless measure *γ* in the tensor form 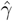 is a multiplication of several factors: the rigidity of cell boundaries and the deviatoric strain along the thickness direction, 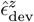. The plane rigidity is system-dependent so we do not consider them in detail here (see Sec. IV for further discussion). The latter term, dependent on the mechanics inside the layer, is our main focus in this section. The tensor form 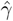 reads:

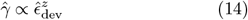

where the in-depth deviatoric strain reads:

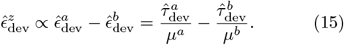

We simply assume that 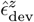 is determined by the difference in the in-plane deviatoric strains 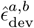 between apical and basal layers, which are 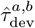 the in-plane deviatoric stresses divided by their respective effective stiffness *µ*^*a,b*^ (in the unit of N/m). Without loss of generality, we can consider the constants of proportionality in Eqs. 14 and 15 to be equal to 1. Then, the magnitude of 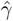 relates to the probability of finding an AB-T1 transition, and the eigenvectors of 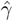 indicate the orientation of the corresponding AB-T1 transition. For the system without shear components, the apical and basal stresses have the same eigenvectors along meridional and circumferential directions, and thus 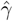 could be reduced to a scalar form *γ* as shown in Eq. 1.

A positive sign of *γ* indicates a tensile strain along the meridional direction with a compressive strain along the circumferential direction, while a negative sign indicates a compressive strain along the meridional direction with a tensile strain along the circumferential direction.

The effective stiffness *µ* depends on how the biopolymers of cell cortex connect, bend, and interact in the lateral sides under the in-plane stresses and it corresponds to the tissue shear modulus resisting the exchange of neighbors along the lateral face. We emphasize that this effective stiffness *µ* is distinct from the cell rigidity controlled by the shape index within the vertex model, because the latter corresponds to dissipative adaptation through cellular rearrangements whereas *µ* is the elastic modulus resisting lateral deviatoric shearing. Generally, the shear modulus of tissues can depend on the pre-compression or pre-expansion of the tissue [40–42] in various ways. Tension-stiffening originates from a bending-to-stretching mode transition, while the mechanism of compression-stiffening originates from jamming [42, 43]. In other cases, a tissue can even display strain-softening due to connections breaking between adherent regions [43].

Supposing that the pre-stress in plane is small, we can establish a phenomenological relationship between the effective stiffness and the pre-stresses in the lowest order as

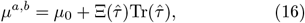

where *µ*_0_ is the intrinsic shear modulus of the material and the trace of stress tensor Tr(*τ* ^*â,b*^) indicates the isotropic tensile or compressive stresses in-plane. The dimensionless coefficient Ξ determines how *µ* depends on the pre-stresses in different kinds of hyperelasticity. Here, we discuss several simple forms for Ξ.

First, we consider linear stress-stiffening, modeled by a positive constant *ξ* for tensile stresses 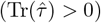 and a negative constant *ξ* for compressive stresses 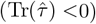; hence Eq. 16 becomes

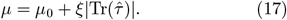

By tuning the value of *ξ*, one can explore varying effects of stress-stiffening in the model. If −*ξ* is set as negative then the model could be extended to a stress-softening case.

If 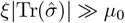 (a strong stress-stiffening), the intrinsic shear modulus can be ignored such that

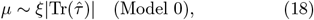

which, normalized by *ξ*, is used for the results in [33]. Oppositely, if 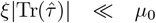 (negligible stress-stiffening), the effective shear modulus is dominated by the intrinsic shear modulus such that

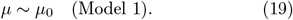

There are other simple forms for *µ* such as: only tension-stiffening;

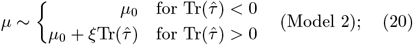

or only compression-stiffening:

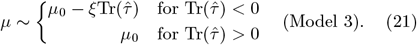

More complex forms can also be expressed, such as strong tension-stiffening while compression-softening:

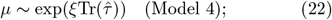

The difference of outputs among these models with the existence of lateral compartments is elaborated in Sec. III E.

### C. In-depth deviatoric strain without lateral load

In this section, we show the analysis of the deviatoric strain profile and the correspondent AB-T1 tendency under the only external loads normal to the surface. This is biologically relevant to a case where the apical or basal shell is subjected to the hydrostatic pressure inside the cells without much tensions along the lateral boundaries.

Substituting *σ*_*T*_ = 0, *σ*_*N*_ = *P* into the force balance equations Eqs. 13, we arrive at:

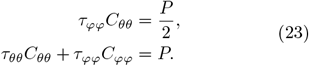

Solving Eq. 23 yields the following relations:

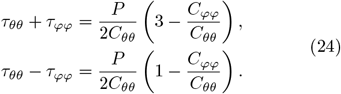

Substituting Eq. 24 into the expression of the in-depth deviatoric strain (Eq. 15), with the *µ* defined in its general form for stress-stiffening (Eq. 17), we obtain the following analytical expression of the in-plane deviatoric strain for apical or basal side as:

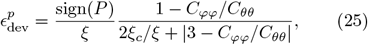

where *ξ*_*c*_ = *C*_*θθ*_*µ*_0_*/* |*P*|.

Note that for an axisymmetric system, *C*_*θθ*_ is always positive, *i*.*e*. the small arc along the circumference is always convex to the polar axis, while *C*_*φφ*_ can be positive for a convex meridian or negative for a concave one, with respect to the polar axis. Fig. 3 is a graphical representation of a normalized deviatoric strain 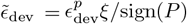 against *C*_*φφ*_ */C*_*θθ*_. When *C*_*φφ*_ */C*_*θθ*_ = 1, the two prime curvatures of a local surface are the same and 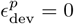, independent of *ξ*.

**FIG. 3.**
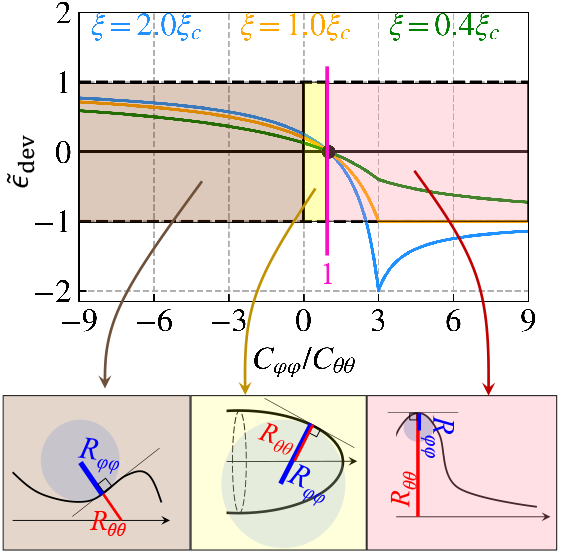
The normalized deviatoric strain 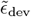 in relation to the ratio of prime curvatures *C*_*φφ*_*/C*_*θθ*_, with various *ξ*. At isotropic curvature condition, *C*_*φφ*_ = *C*_*θθ*_ (magenta line). Bottom panel exhibits three typical examples of different curvature ratio: (left) brown for *C*_*φφ*_*/C*_*θθ*_ *<* 0; (middle) yellow for 0 *< C*_*φφ*_*/C*_*θθ*_ *<* 1; (right) pink for *C*_*φφ*_*/C*_*θθ*_ *>* 1. The arrow indicates the polar axis. The radius of curvature *R*_*θθ*_ = 1*/C*_*θθ*_ and *R*_*φφ*_ = 1*/C*_*φφ*_ are highlighted by the red and blue lines respectively.

When *ξ* ≫ *ξ*_*c*_ (strong stress-stiffening, blue curves in Fig. 3), the largest magnitude of 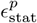 occurs at *C*_*φφ*_ */C*_*θθ*_ = 3 with its value

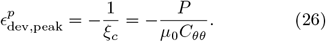

For a shape elongated along the polar axis without bumps, *C*_*φφ*_*/C*_*θθ*_ = 3 is not feasible. In this case, the magnitude of *ϵ* increases when *C*_*φφ*_*/C*_*θθ*_ approaches −∞, where the curvature anisotropy |*C*_*φφ*_ −*C*_*θθ*_| becomes largest.

When 0 *< ξ* ≪ *ξ*_*c*_ (weak stress-stiffening, green curves in Fig. 3), the magnitude of 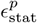 always increases with anisotropy of curvature. The largest magnitude of deviatoric strain occurs with value 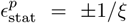 when *C*_*φφ*_*/C*_*θθ*_ → ∓∞.

When *ξ* ~ *ξ*_*c*_ = *µ*_0_*C*_*θθ*_*/*|*P* | (orange curve in Fig. 3), the largest magnitude of 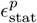 occurs where *C* _*φφ*_ */C*_*θθ*_ *>* 3. Large *C* _*φφ*_ */C*_*θθ*_ corresponds to geometries such as bumps, see the right bottom panel in Fig. 3.

Although the strength of stress-stiffening (value of *ξ*) affects the magnitude of deviatoric strain in different ways for *C*_*ϕϕ*_*/C*_*θθ*_ *>* 1, the behaviors of deviatoric strain are robust against *ξ* for the region *C*_*ϕϕ*_*/C*_*θθ*_ *<* 1. With *C*_*ϕϕ*_*/C*_*θθ*_ *<* 1, Eq. 25 reduces to:

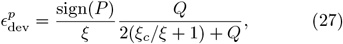

where *Q* = 1 −*C*_*φφ*_*/C*_*θθ*_ *>* 0. For a modulus model *µ* with stress-stiffening (*ξ >* 0), the largest magnitude of deviatoric strain always occurs at the largest *Q, i*.*e*., smallest value of *C* _*φφ*_ */C*_*θθ*_.

We further narrow the cases to only convex surfaces, *i*.*e. C* _*φφ*_ ≥ 0 (the yellow region, or the middle bottom panel in Fig. 3). This curvature regime is typical for a regular elongated axisymmetric shape, such as an ellipsoid or cylindrical tube. Then the largest magnitude of deviatoric strain occurs at *C* _*φφ*_ = 0. This corresponds to the trunk region of a prolate ellipsoid. For a modulus model of stress-softening (*ξ <* 0), the right hand-side of Eq. 27 has a singularity at *Q* = − 2(*ξ*_*c*_*/ξ* + 1), dependent on the the value of *ξ*, and *C*_*θθ*_ and *µ*_0_ as well as the hydrostatic pressure *P*.

Next, we analyze the dependency of AB-T1 transition tendency *γ* on the curvature ratio. Let *c* denote the ratio between two principal curvatures *C* _*φφ*_ */C*_*θθ*_. In the limit where hydrostatic forces dominate, the AB-T1 tendency profile *γ* as a function of the arc position *s* becomes

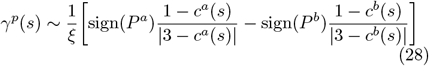

for *ξ/ξ*_*c*_ ≫ 1.

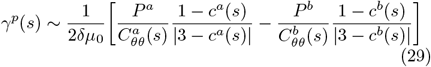

Since the cell height *e* is much smaller than the radius of curvature (our model assumption),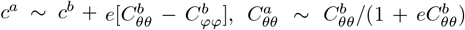, then the profile of *γ*^*p*^(*s*) is approximately proportional to the normalized hydrostatic deviatoric strain at the basal side 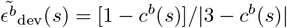 as

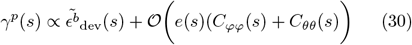

with the linear coefficients determined by the hydrostatic loading *P* ^*a,b*^ and a negligibly small correction from the cell height. Hence, the AB-T1 tendency under hydrostatic conditions, *γ*^*p*^, approximately follows the profile of in-plane deviatoric strain, which means *γ* is near zero at locations with isotropic curvature and maximizes at the locations with largest curvature anisotropy for an elongated axisymmetric shape.

### D. In-depth deviatoric strain with lateral load

For the calculations under hydrostatic pressure, we have ignored the existence of the lateral membranes. In order to analyze the contribution from the lateral side, we first consider how lateral membranes rest on the curved surface. We first approach this problem by considering the free energy of the system in a discrete description of cell packing. The cell packing - which determines the cell areas on apical, basal and lateral sides - reaches a stable configuration when the system finds its minimal free energy. Following the established literature of vertex models [44–46], we describe the forces regulating the cell shape tissue as a derivative of the following free energy function:

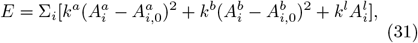

where 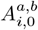 is the preferred cell area at the apical (or basal) layer for each cell *i. k*^*a,b*^ is the apical (or basal) elasticity coefficient and *k*^*l*^ is the lateral tension strength; 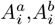 and 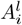 are the areas of cell *i* at apical, basal and lateral surfaces respectively.

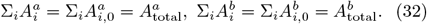

The packing equilibrium corresponds to the minimum of the free energy (Eq. 31) under the surface constraints given by Eq. 32.

If the lateral membrane is far less contractile than the apical and basal membranes, *i*.*e*.,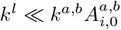, the minimization of this free energy will cause the cell to optimize its area towards 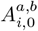 in the apical and basal sides and the lateral membrane will tilt when the local curvatures of the shell change along the surface. By contrast, if 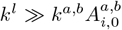, the minimization of the free energy leads to the lateral membrane orientating perpendicular to the apical and basal sides, with the cell apical area becoming a normal projection of the basal area, depending on the local curvatures. This can be seen from calculating the functional derivatives of Eq. 31 as below.

For a demonstrative purpose, we show a derivation in a 2D equivalent (meridional cross-section as shown in Fig. 2b) and assume the preferred area of cells is homogeneous along the surface such that 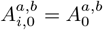. We first discuss a case without the single-cell volume constraint and then extend to a case with the volume constraint. The free energy in a 2D system is

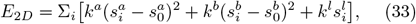

where 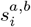 are the arc lengths of the cell at the apical or basal sides and 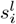 is the length of the cell lateral membrane and 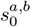 are the preferred cell lengths at the apical or basal side. Note that cell height 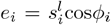, where *ϕ*_*i*_ is the tilt of lateral membrane of cell *i*. Minimizing this free energy constrains the cell shape so that the functional derivatives of the free energy become zero:

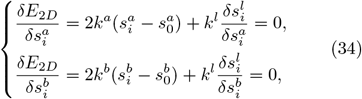

where 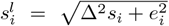, depicted by the line *ss* ′ in Fig. 2b. The surface constraint (Eq. 32) accordingly turns into a 1D form as

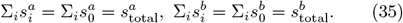

For simplicity, we assume *k*^*b*^→ ∞ (due to the symmetry of the energy function, this assumption is equivalent to the case with a finite *k*^*b*^ but *k*^*a*^→ ∞), meaning that the basal layer is solid and thus the second equation in Eq. 34 can be ignored with 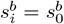.

Let Δ*s*_*i*_ denote the difference between the 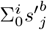, which is the accumulated normal projection length from the basal arc, and 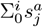, which is the accumulated apical arc length at cell *i*. Let *κ*_*i*_(*C*_*i*_, *e*_*i*_) be the normal projection rate merely depending on the curvature *C*_*i*_ and cell height *e*_*i*_ of cell *i*, and 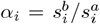 be the ratio of basal length over apical length of cell *i*, then

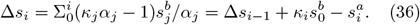

Eq. 34 then becomes

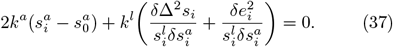

Without the volume constraint, the cell height distribution *e*_*i*_ is a consequence of multiple regulators, hence for simplicity we just take 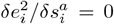. Then, Eq. 37 reduces to:

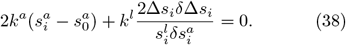

Note that the ratio *α*_*i*_ varies from 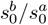 to 1*/κ*_*i*_ as the lateral stiffness changes. If lateral contractility does not exist, *i*.*e*., *k*^*l*^ = 0, then from Eq. 38, 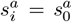 and 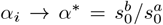, which is constant along the surface. Thus, from Eq.36, Δ*s*_*i*_ becomes

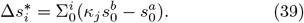

On the contrary, if apical contractility is very small, *i*.*e*., *k*^*a*^ ≈ 0, from Eq. 38, we have Δ*s*_*i*_ = 0. From Eq. 36, it then follows *α*_*i*_ = 1*/κ*_*i*_, which is determined by the layer curvature *C*_*i*_ and height *e*_*i*_.

For general cases with non-zero *k*^*l*^ and *k*^*a*^, inserting Eq. 36 into Eq. 38, we have

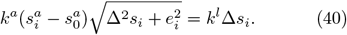

Let 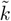 be *k*^*l*^*/k*^*a*^*e*_*i*_ as a ratio of lateral contractility over plane rigidity. For a limit 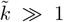, *i*.*e*., rigid lateral contractility dominates over the plane rigidity, we get

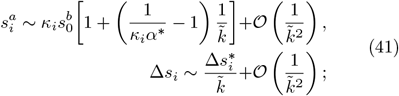

Hence, the corresponding tilt angle is

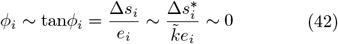

which means the lateral boundaries are perpendicular to the layer surface.

For another limit 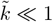, *i*.*e*., weak lateral contractility as compared to the plane rigidity, we get

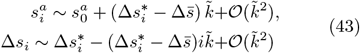

where 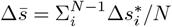.

We assume for Δ*s*_*i*_ *≪ e*_*i*_, the corresponding tilt angle *ϕ* of the lateral membrane is approximated as tan*ϕ*_*i*_ = Δ*s/e*_*i*_ and for the limit of zero lateral contractility,

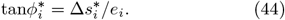

In both the rigid lateral (Eqs. 41) and soft lateral (Eqs. 43) cases, Δ*s*_*i*_ is a small deviation from the value at the extreme end. Given the importance of Δ*s*^∗^, in the following section we present a continuum expression of Δ*s*^∗^ with the corresponding tilt angle *ϕ*^∗^ and discuss its relation to surface curvature under more general assumptions.

#### 1. A continuum model for zero lateral-contractility limit

Fig. 2b illustrates a meridional cross section for an arbitrary axisymmetic shell. A surface element *dA*^*b*^(*φ, θ*) located at the arc position *s*^*b*^ on the basal side (grey curve), can be mapped to another surface element *dA*^*a*^(*φ*′, *θ*^l^) located at the arc position *s*^l^ in such a way that the accumulated number of cells from the head of the object to *s* on the basal side is the same as the cell number accumulated from the head to *s*^l^ at the apical side. On both surfaces, the surface element could be represented by the small arc *ds* as *dA* = *r*(*s*)*dθds*, where *r*(*s*) is the distance to the polar axis. Note that *r*(*s*)*dθ* and *ds* are along two orthogonal directions, the circumferential and meridional directions, respectively. For the simplicity of notation, we just discard the superscript *b* for geometrical quantities on the basal side in the derivation.

Given our assumption of an axisymmetric surface, the 2D integral of surface element *dA* over the whole shell surface can be reduced to a 1D integral with only the meridional variable from 0 to *s*. The accumulated number of cells *N* from the head to *s* on the basal side is given by

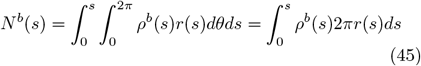

and it is equal to the accumulated number of cells *N*^*a*^ on the apical side:

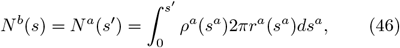

where *ρ*^*b*^(*s*) or *ρ*^*a*^(*s*) is the cell density on the basal or apical surface, The density *ρ*^*a,b*^(*s*) is determined by minimizing the membrane tensions on apical, basal and lateral sides. Here, we do not consider any other cues guiding cell location within the tissue environment.

The cell density at the apical side is related to that at the basal side at *s* as *ρ*^*a*^(*s*) = *α*(*s*)*ρ*^*b*^(*s*). Since the total number of cells are the same at the two sides, the distribution of apico-to-basal ratio of density *α*(*s*) must follow:

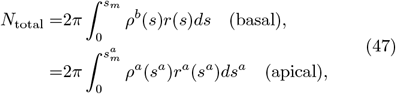

where the integration of *ds*^*a*^ (apical) or *ds* (basal) is over the whole meridional range *φ* ∈ [0, *π*] and 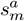 or *s*_*m*_ represents the half meridian and *N*_total_ is the total cell number covering the shell. When the apical element *dA*^*a*^ is only a normal projection of the basal element *dA* with a small distance *e*(*s*), then one can obtain:

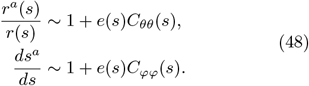

Inserting Eq. 48 into Eq. 46 leads to

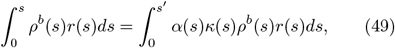

where

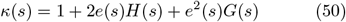

is the normal projection ratio, *H*(*s*) is the mean curvature

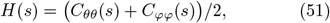

and *G*(*s*) is the Gaussian curvature

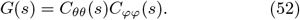

Equation 49 now only depends on quantities with superscript *b*, so for neatness we omit this superscript from here and write *ρ*^*b*^(*s*) = *ρ*(*s*), *N*^*b*^(*s*) = *N* (*s*). Reorganizing the integration on the right hand side, we can transform Eq. 49 into

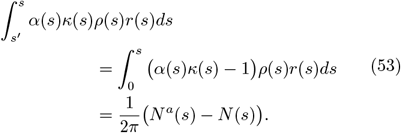

For *s*′ − *s*→ 0, the left hand side of Eq. 53 can be approximated by Δ*s*(*s*), the difference in arc length between *s* and *s*′, as

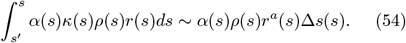

Accordingly, from Eq. 53 we get

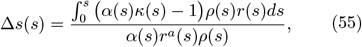

and then derive the tilt angle *ϕ* as:

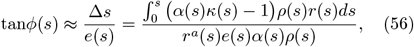

or in a more compact form

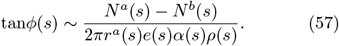

As we discussed in the previous section, different lateral contractility induces different values of ratio *α*(*s*). For the strong lateral contractility limit, *α*(*s*) → 1*/κ*(*s*) so that both Δ*s*(*s*) and *ϕ*(*s*) become zero. For the zero lateral contractility here, *α* is independent of the local curvature and becomes the ratio of the two meridians 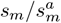:

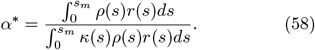

Inserting Eq. 58 into Eq. 56 gives

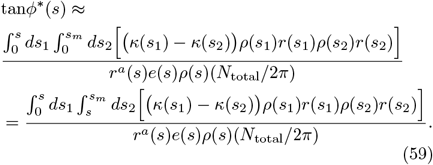

As the cell size is much smaller than the radius of curvature, *κ*(*s*_1_) − *κ*(*s*_2_) ~ 2[*e*(*s*_1_)*H*(*s*_1_) − *e*(*s*_2_)*H*(*s*_2_)] with the second order term neglected. To clearly see the dependency of *ϕ*^∗^ on curvature, we transform the integration of *ds* in Eq. 59 into integration by local cell number *dN* (*s*) = 2*πρ*(*s*)*r*(*s*)*ds* as

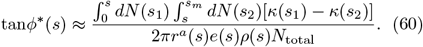

The integral 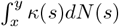 could be alternatively expressed as 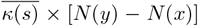, where the notation ∗ is defined as

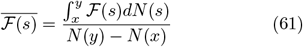

is the weighted average of any quantity ℱ (*s*) in the range of *x < s < y*. Hence, Eq. 60 be expressed as

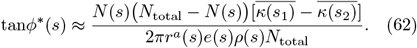

As long as the change of cell height *e*(*s*) with *s* is smaller than the change of curvature, we can approximate the difference of *κ* mainly by the change of mean curvature as 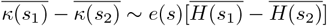.

We can relate the difference between the weighted average of the mean curvature to the mean curvature gradient. For 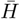 averaged from *x < s < y*, according to the integral mean value theorem, we can always find an 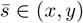 such that 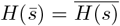. Hence, the difference of 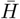 can be re-expressed as

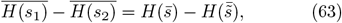

where 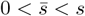 and 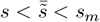. Since 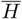 is the average weighted by the cell number at 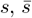 and 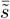 should be close to where the local cell number *dN* (*s*) is large, *i*.*e*., where *ρ*(*s*)*r*(*s*) is large. If the cell density does not radically change with *s, r*(*s*) will dominate where 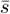 and 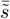 locate.

When *s* → 0 (head) or *s* → *s*_*m*_ (tail), *i*.*e*.,, at the two polar ends, *ϕ*^∗^ is close to zero because *N* (*s*) *N*_total_ − *N* (*s*)) → 0 and the contribution from the curvature becomes trivial. When *s* is neither close to the head nor the tail, if the surface is convex, *r*(*s*) is large and 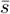 and 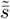 will be in a vicinity of *s*, where the distance 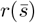 and 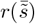 is relatively large. According to the mean value theorem, we could find another 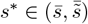 such that

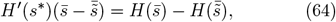

where *H*′ is the gradient of curvature and *s*^∗^ is even closer to *s* than 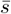 and 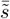. If the gradient of curvature is also continuous and differentiable (as seen in the ellipsoidal or tubular structures in biological systems), *H*′ (*s*^∗^) ≈ *H*′ (*s*) + (*s*^∗^ − *s*)*H*^″^(*s*); therefore, given a steep mean curvature gradient at *s*, we predict in the zero-lateral-contractlity limit that there is a large tilt angle *ϕ*^∗^(*s*), as long as *s* is not at the head or the tail. Additionally, since 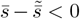, a negative gradient of curvature along *s* corresponds to the positive tilt angle towards the head. In other words, the lateral tilt will lean towards the position of higher positive curvature.

#### 2. Inhomogeneous layer height and density

As shown in Eq. 60, the tilt with the zero lateral contractility in general cases is related not only to the curvatures, but also to the cell height *e*(*s*) and cell density *ρ*(*s*) along the surface. Note that *e*(*s*)*/ρ*(*s*) is proportional to the cell volume profile along the surface and a constant volume constraint corresponds to the condition that *e*(*s*) ∝ *ρ*(*s*). In this section we analyze the contributions from the inhomogeneous cell height and density profile to the lateral tilt profile.

Using the mean value theorem to eliminate the integral in Eq. 59, we obtain:

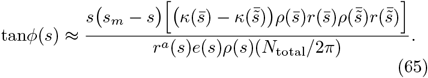

By setting a homogeneous density *ρ*(*s*) ~ *ρ*_0_ and *e*(*s*) ~ *ε*, we arrive at a tilt profile purely depending on the geometry of the surface:

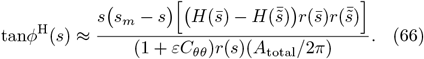

We now evaluate the contributions from height modulation and basal density modulation separately. We define *ŝ* such that the total cell number at the apical side *N*_total_ = *ρ*(*ŝ*)*A*_total_, where *ρ*(*ŝ*) is a weighted average of density from *s* = 0 to *s* = *s*_*m*_. According to Eq. 65, the tilt profile with modulated inhomogeneous density becomes

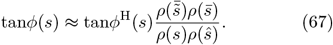

If *ρ*(*s*) is nearly homogeneous as |*dρ/ds*| ≪ 1, we assume *ρ*(*s*) ~ *ρ*_0_[1 + *η*(*s*)(*s* − *ŝ*)*/s*_*m*_] with |*η*(*s*)| ≪ 1. Then,

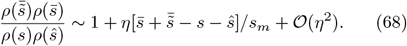

Recall that *ŝ* is the averaged position weighted by *ρ*(*s*)*r*(*s*) while 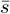 and 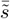 are the averaged positions weighted by *κ*(*s*)*ρ*(*s*)*r*(*s*). With a radius of surface curvature much larger than the typical cell size (an assumption of our model) - *κ* is only slightly larger than 1 - we approximately have 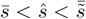. In this case, the first order term in *η* is negligible. In particular,

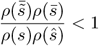

is valid when *ρ*(*s*) is a monotonic function. Therefore, the tilt angle under mild inhomogeneity of density is always slightly smaller than the scenario with homogeneous density.

The tilt profile with modulated inhomogeneous cell height *e*(*s*) can be evaluated similarly, supposing *e*(*s*) = *ε*(1 + *η*′(*s*)(*s* − *ŝ*)*/s*_*m*_) with |*η*′(*s*)| ≪ 1. Clearly, the value of *ε* has negligible effect on the result as long as *εH*(*s*) ≪ 1 (our basic model assumption) is valid.

Then, the tilt profile corrected by inhomogeneous cell height is

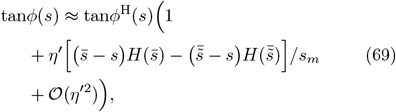

which has a more significant first order correction term in 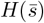 than Eq. 67. Therefore, inhomogeneity in cell height causes greater deviation of the tilt angle from the homogeneous limit *ϕ*^H^ than inhomogeneity in cell density. In Fig. 4, we show the tilt profile and corresponding phase diagram for the AB-T1 transition measure for a prolate ellipsoidal system with aspect ratio *r*_2_*/r*_1_ = 0.4 and *ε/r*_1_ = 0.05; *r*_1_, *r*_2_ are the half major (or minor) axis lengths. The horizontal axis is the relative distance to the head and *d* = 1 represents the trunk. Tissue height and basal density are modulated linearly with *s*, with coefficients *β* and *λ* respectively. Modulation of density slightly suppresses the final tilt angle (the continuous curves slightly lower than the dashed curves). Meanwhile, modulation of height (varying *β*) affects the tilt more significantly, not only affecting the magnitude but also the shape of the distribution profile (as compared with the yellow lines, which corresponds to a homogeneous or zero modulation limit).

**FIG. 4.**
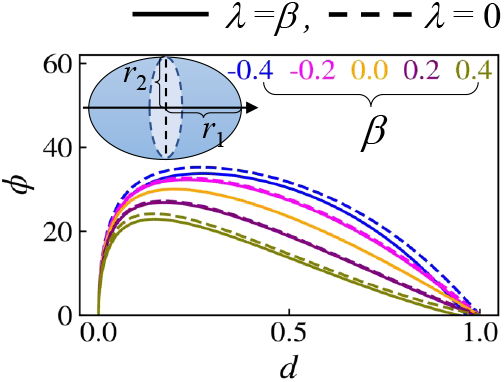
Results with an inhomogeneous basal density distribution modulated as *ρ*(*s*) = *ρ*_0_ [1 + *λ*(*s/s*_1*/*4_ − 1*/*2)] and inhomogeneous cell height distribution *e*(*s*) = [*ε* 1+*β*(*s/s*_1*/*4_ − 1*/*2)], where *s* is the arc length along the prime meridian surface and *s*_1*/*4_ is the 1*/*4 arc length.The system is a prolate with aspect ratio *r*_2_*/r*_1_ = 0.4 and cell height *ε/r*_1_ = 0.05. A comparison of tilt angle profile between *λ* = *β* (continuous) and *λ* = 0 (dashed) for varying *β* from −0.4 to 0.4.

In conclusion, assuming the change in cell height and cell density are relatively small to the change in curvature along the surface, the generalized cell tilt profile is more sensitive to cell height inhomogeneity than cell density. Hence, it is a good approximation to keep density homogeneous and consider only cell height modulation, when considering the onset of cell tilt under curvature constraints.

### E. AB-T1 transitions under different material properties

Having verified that *γ* is a good approximation (at least within experimentally plausible parameter regimes) of the AB-T1 transition rate, we can now explore how different material properties alter the behavior of cell rearrangements under different tissue curvatures. Before introducing the outputs of different models, we first recall the baseline results obtained with model 0, which correspond to the main results in Ref. [33]. The shell geometry is simply set as a prolate ellipsoid with the major half length *r*_1_ and minor half length *r*_2_ giving an aspect ratio *r*_2_*/r*_1_ = 0.4. The external load acts through differences in pressures ΔΠ^*a*^ and ΔΠ^*b*^ on the apical and basal sides respectively, together with the lateral tension *T* (Fig. 5a). We define a measure for the maximum probability of AB-T1 transitions, which we refer to as the significance of the peak of |*γ*|:

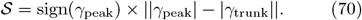

**FIG. 5.**
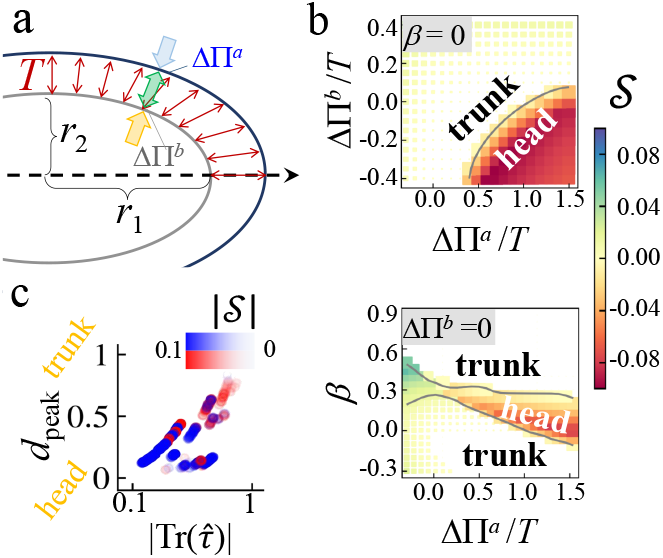
Output of Model 0 in the zero lateral contractility limit with a prolate geometry. (a) External forces acting on the bilayer in a curved environment. (b) Phase diagram of with ΔΠ^*a*^*/T* and Δ^*b*^*/T* at *β* = 0 (top), and ΔΠ^*a*^*/T* and *β* at ΔΠ^*b*^ = 0 (bottom). The color represents the significance of the peak 𝒮 (Eq.70) and the size of the data point size scales as ∝ (1 − *d*_peak_) ^2^, the squared distance of the peak to the trunk. The position of peak of |γ|, shifts from head to trunk with the increase in the isotropic stress in the layer 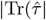. Blue color corresponds to *γ*_peak_ *>* 0 and red to *γ*_peak_ *<* 0. The opaqueness of the color represents |𝒮| the magnitude of significance of peak.

This quantity is non-zero only if the peak is prominent and far away from the trunk.

The phase diagram of Model 0 is presented in Fig. 5b (a 3D version was also presented in [33] Figs. 3 and 4), where the maximum probability of AB-T1 transitions occurs either at the trunk or head of the prolate ellipsoid. The transition between these two cases depends on pressure differences compared to the lateral tension at apical and basal layers, and the inhomogeneous layer thickness. The size of the blocks in Fig. 5b indicate the squared distance of the *γ*_peak_ to the trunk *d*_peak_. From Fig. 5c, it can be observed that *d*_*peak*_, the position of peak tendency of AB-T1 transitions is roughly proportional to the magnitude of in-plane isotropic tension 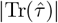, *i*.*e*., a smaller isotropic stress leading to a smaller lateral shear modulus induces AB-T1 transitions mostly at the head of the prolate.

Next, we show the comparison of *γ* under the five previous highlighted effective shear modulus. In Models 2, 3, 4, we set *µ*_0_ = *ξ* and all the *γ* shown here are normalized by *r*_1_*/δξ*, where *r*_1_ is the half of the major axis of the prolate ellisoid.In Fig. 6, we show the comparison of the spatial profile of *γ* across different lateral shear modulus models (see Eqs.18-22) with the black dots highlighting the peak of |*γ*|. The corresponding phase diagrams for Models 2-4 defined in the parameter space of ΔΠ^*b*^*/T* −ΔΠ^*a*^*/T* and *β* −ΔΠ^*a*^*/T* are presented in Fig. 7. We can see for different models of the effective modulus *µ*, the phase diagrams of significance of peak have different boundaries between the trunk region (white) and the head region (colored) in the parameter space. Both material properties and tissue geometry play an important role in the occurrence and positioning of the AB-T1 transitions. Different models may be relevant depending on specific experimental conditions.

**FIG. 6.**
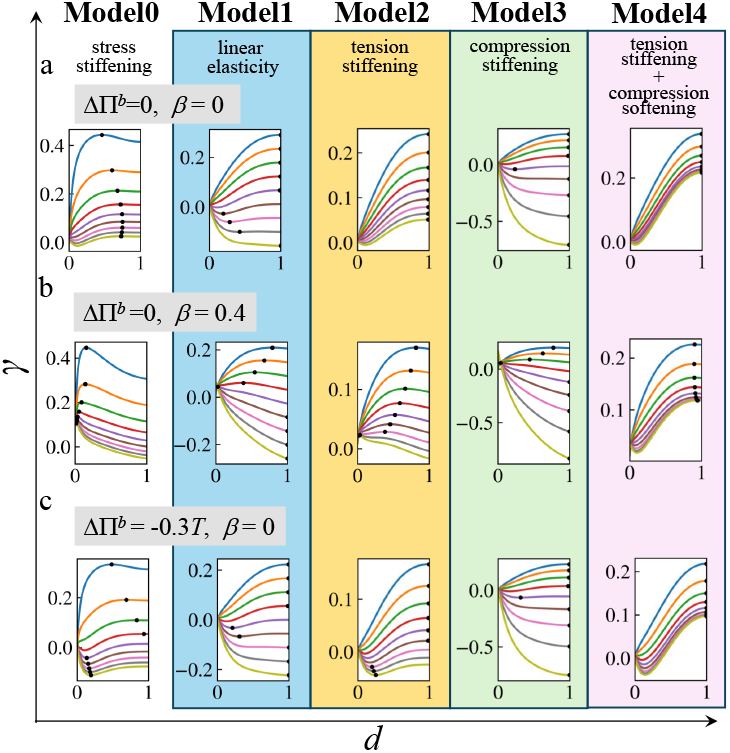
Comparison among different models for effective shear modulus for a prolate ellipsoid *r*_2_*/r*_1_ = 0.4: Distribution of AB-T1 transition tendency *γ* for varying apical pressure differences ΔΠ^*a*^*/T* from top to bottom: −0.3 (skyblue), 0 (orange), 0.3 (green), 0.6 (red), 0.9 (purple), 1.2 (brown), 1.5 (pink), 1.8 (grey), 2.1 (golden), with three different value sets of the basal pressure difference ΔΠ^*b*^ and tissue height inhomogeneity *β* (panel a-c). Horizontal axis *d* is the relative distance; *d* = 0 indicates the head of a prolate ellipsoid and *d* = 1 the trunk. Black dots indicate the peak of |*γ*|.

**FIG. 7.**
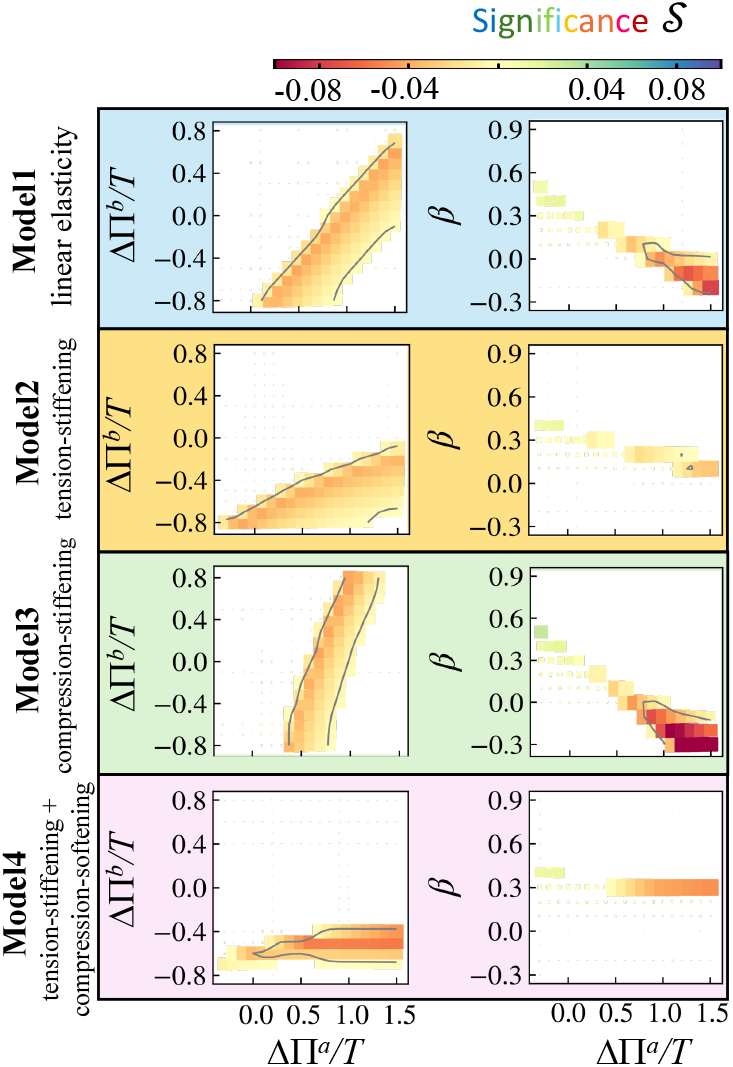
Comparison among different models of effective shear modulus for a prolate ellipsoid *r*_2_*/r*_1_ = 0.4. Diagrams of the significance of the peak of *γ*, which is defined as sign(*γ*_peak_) × ||*γ*_peak_| − | *γ*_trunk_||. From top to bottom: Model 1, Model 2, Model 3, Model 4. White regions indicate peak of |*γ*| at trunk, whereas coloured squares indicate peak of |*γ*| at near the head. The corresponding diagrams for Model 0 are presented in Fig.5b and [33] Fig.3 and 4.

## IV. DISCUSSIONS

Here, we have focused on the differences of stress/strain anisotropy in the apical and basal layers in a cell monolayer. We have interpreted these results in terms of the apico-basal T1 transition rate and shown this is consistent with numerical simulations. Our theory is based on continuum mechanics and does not assume specific properties for the apical and basal sides in each single cell. This means that the theory itself should be applicable to other bi-layer systems in which those layers have different directionalities of stress/strain anisotropy from one another, just by modifying specific assumptions.

Technically, one can apply any external force field and select proper assumptions on the mechanical properties according to the specific systems of interest and solve the equation Eq. 9 (or Eq. 13 if it is an axisymmetric system). In this paper we did not discuss any external forces originated in plane, as the main focus was on the treatment of forces from the lateral structure between two layers. However, our theoretical framework allows for any local in-plane forces to be added to the force balance equation. These forces could be generated during active cellular processes and may be necessary to consider in other tissue mechanical processes.

There are several other details in our theory that depend on the system of interest. As an example of such system-dependent assumptions, here we investigated the difference between Models 0-4 for the effective shear modulus. For another example, the form of the potential function (Eq. 31) is system-dependent. In particular, the lateral tension part, or how the force is transmitted between the two layers, can be differ from system-to-system. Besides the mathematical potential form of the lateral tension, the difference between the strong and weak lateral-tension limit may be important; here we focused on the weak limit which will give rise (Subsec. III D 1).

Beyond external forces, material properties such as the plasticity or fluidity of cell boundaries (which is controlled by the shape index in the vertex model as discussed in Sec. II and Fig. 1b), or the non-linearity in the lateral elastic modulus (as discussed in Sec. III B) are also system-dependent and can impact the AB-T1 transition rate. In particular, a higher lateral plasticity might decrease the occurrence of AB-T1 transitions (Fig.1c) by reconciling the in-depth anisotropic stresses through the floppy adaptation of lateral interface positioning.

Our theory is appropriate to describe particular *in vivo* systems. In *Hydra* [47, 48], a bilayer cell sheet forms, consisting of the endodermal and ectodermal layers respectively. The actin fiber directions in each sheet are perpendicular to each other [47, 48]. We predict that there will be differences in stress anisotropy between endodermal and ectodermal actin cytoskelstons leading to potential AB-T1 transitions. Another example is the crisscross multilayer of myoblasts [49]. Since the principal axes of a stress tensor are expected to align with the major/minor axes of the elongated cell shape, a similar anisotropy in the stress between layers may be expected there. Testing of models is achievable through live imaging and semi-automated image analysis [29, 50, 51]. It will be interesting to study how the measure *γ*, which we have introduced in this paper and [33] is correlated with observable phenomena in such systems.

## V. CONCLUSIONS

Layers with lateral connections are common in biological systems. Here, we have provided detailed analysis of a continuum framework to describe mechanical forces within a 3D curved layer supported by a lateral mesh, substantially building on [33]. We focused on the apical and basal layers of an epithelial monolayer as an example (Sec. II) and scrutinized how layer curvature and cell mechanics interplay to create stress anisotropy, potentially resulting in irregular 3D cell shapes including AB-T1 transition, which is characterized by scutoid-like geometries (Sec. III). Such detailed consideration clarified the specific assumptions used in our theory based on the general continuum framework. We then discussed the capability of this framework to be applied to broader biological systems, including clarifying the system-dependent assumptions (Sect. IV). We expect that this framework can serve to explain the morphological features in various multilayered biological systems with distorted stress/strain anisotropy.

## VI. ACKNOWLEDGMENTS

We thank Jacques Prost for discussions that motivated this work. Y. L is funded by start-up support from Fudan University. T.H. is funded by a Mechanobiology Institute seed grant. T.E.S. and S.T. were funded by EPSRC Physics of Life grant (R.MRCB.1137) and start-up support from the University of Warwick. T.E.S. and S.T. are grateful to support from grants from the NSF (PHY-2309135 to the Kavli Institute for Theoretical Physics (KITP) and PHY-2309135) and the Gordon and Betty Moore Foundation (Grant No. 2919.02) which supported their stay at KITP, where part of this work was completed.

